# Giant pore formation in vesicles under msPEF-induced electroporation: role of charging time and waveform

**DOI:** 10.1101/2024.06.14.598990

**Authors:** Nalinikanta Behera, Rochish M. Thaokar

## Abstract

Giant unilamellar vesicle is the closest possible prototypical model for investigating membrane electrodeformation and electroporation in biological cells. In this paper, the effect of membrane charging time on vesicle electroporation, an unresolved issue, is exclusively investigated under milli-second pulsed-electric-field (msPEF) of different waveforms, using numerical simulations. The existing analytical models uncover several fundamental features of cell or vesicle electroporation, but remain far from being realistic as electrodeformation effects were neglected. Our numerical approach, which implements the effect of electric stretching on membrane tension and precise calculation of pore energy, successfully predicts the formation of giant pores of *O*(*µm*) size as observed in past experiments. The poration zone is found to extend up to certain angles from the poles, termed critical angles. Increase in charging time delays pore formation, decreases the pore density as well as trim downs the poration zone. Counterintuitively, this effect promotes significant pore growth. Thus, more pores evolve into giant pores resulting in higher fractional pore area. Moreover, there exists a cut-off charging time above which pore formation is completely inhibited. This phenomenon is particularly pronounced with bipolar pulses. Comparisons with the past experimental results reveal that electrodeformation-poration-induced membrane surface area variation and that induced by electroporation evolves in a similar fashion. Therefore, although the agreements are qualitative, the present electroporation model can be used as the simplest tool to predict the transient electrodeformation of a porated vesicle in the laboratory experiments.

## Introduction

Electroporation refers to the electric field-induced formation of aqueous pores in cell membranes, which enables electrodiffusive species transport between intracellular fluid (or cytoplasm) and extracellular fluid [25, 16]. In the rapidly evolving landscape of biomedical research, the significance of electroporation as a cutting-edge therapeutic method cannot be overstated. Among the nonviral transfection methods, electroporation is considered as an inexpensive research tool for delivering interfering RNAs into cells to selectively regulate gene expression [6]. This method is useful for probing the mechanism of death of bacterial-infected cells and immunosuppression in the tumor microenvironment [5, 35]. Both in vitro and in vivo electroporation-based chemotherapy or electrochemotherapy (ECT) are widely used for promoting the uptake of genetically engineered cells and chemotherapeutic drugs into cells [4].

The transmembrane potential (TMP) is the key parameter dictating electroporation, refers to the voltage difference across the membrane. If the TMP crosses a threshold limit (*≈* 1*V*), the permeability of the membrane considerably increases attributed to molecular rearrangement, forming hydrophilic passages (or pores) of *O*(10^*−*9^)m [1, 10, 8]. However, understanding electroporation, even using the state-of-art imaging tools is challenging due to the enormous number of short-lived nanometer sized pores. The alternative approach of quantifying electroporation is measuring the change in concentration of exogenous substances or the leakage of cellular substances. There is currently no established experimental tool for delivering the spatial and quantitative aspects of electroporation. On the other hand, in the last two decades, computational studies employing either molecular dynamics or continuum approaches have elucidated the underlying physics of electroporation to some extent. However, molecular dynamic simulations cannot resolve the spatio-temporal pore formation on a length scale greater than a few 10s of nanometers. Consequently, continuum models have gained widespread acceptance within the research community, providing an understanding of diverse aspects such as spatiotemporal variations in pore number, its size and TMP.

During the early stage of theoretical developments, a substantial amount of effort was devoted to formulating the membrane pore energy [1, 38, 2]. The major drawback with these theories was that the pore radius must be very small compared to the membrane thickness (*h ≈* 5nm). These are, therefore, valid for ultra-short weak pulses, wherein the pore growth is negligible. However, most electric pulses of interest in electroporation are a few hundred microseconds and strong enough to create much greater pore sizes up to 1*µm* or greater. Krassowska and group addressed this limitation of the earlier models in a series of theoretical works [22, 3, 21, 24, 36]. Neu and Krassowska [24] suggested formulations for electric energy for large pores, using which, in a subsequent study, Smith *et al*. [36] were able to predict the growth of small pores into big pores in a uniformly polarized membrane. Their initial study on the local analysis of a planar membrane could predict highly non-linear pore formation and growth and the simultaneous coupled variation of the transmembrane potential. The physics of electroporation on a cellular level can be much more complex than those of a local small planar patch of the membrane, predominantly on account of spatial variation of the electric field [26, 7, 27, 18]. Using computational simulations, a number of studies demonstrated that for spherical biological cells, the TMP, pore number, and pore size are nonlinearly coupled, resulting in nonintuitive transient variations in TMP along the membrane [17, 14, 11, 34, 13].

Despite the numerous studies, there are two crucial facets that remain poorly addressed (a) the pore growth, which results in very large pores (of the order of 100s of nm), and (b) the influence of membrane charging time (*τ*_*c*_) on this process, and this forms the basis of the present work. Controlling the pore growth is the utmost concern in surgical procedures as it controls the reversibility of cell transformations. However, the time period over which the transition from reversible to irreversible electroporation occurs is quite sensitive to membrane charging time as well as cellular conditions. Hence, achieving specific desired electroporation characteristics in a cell or a tissue can be challenging in clinical set-ups. Compounding this challenge is the lack of robust theoretical support for experimental findings, primarily due to the absence of an accurate pore growth model. Pore growth in the reported numerical models was considered only a function of edge tension, membrane tension, and TMP [36, 17, 37, 13]. However, the contribution of the electric tension created by Maxwell’s stress was not taken into account. Consequently, Krassowska and Filev [17], found almost no variations in the average pore size on increasing electric field strength above (*≈* 30*kV/m*). This imposes serious questions on the modeling approach as the membrane clearly tends to stretch more owing to an increase in Maxwell’s stress (*∝ E*^2^), thereby resulting in increased pore size. In fact, in experiments by Riske and Dimova [28], the formation of pores of the size of a few *µ*m is well reported. Whereas the aforementioned numerical models failed to predict the pores of size beyond *O*(10^*−*7^)m, which can be attributed to the negligence of Maxwell’s stress effects. Furthermore, an in-depth literature review suggests that in the context of electroporation, only the cases of very low charging time, i.e., *τ*_*c*_ *≪* 1*µ*s, were examined considering cytoplasmic or extracellular fluids of finite conductivity (*O*(1)*S/m*). For such cases, the charging time is less than nanosecond, that usually leads to rapid but controlled poration. Giant unilamellar vesicles (GUVs), which have been widely used as a biomimic for biological cell, offers control over inner and outer fluid conductivities, and thereby provides an easy way for regulating the membrane charging time. In the earlier experiments conducted to study the combined electrodeformation and electroporation of GUVs [29, 28, 19], the fluid conductivities varied between *O*(10^*−*4^) *− O*(10^*−*3^) S/m. The charging time, accordingly, can reach up to *O*(1)*ms*. Such extended charging time can retard TMP development, and thereby, potentially alter the poration in unconventional ways. Here, the key question is what will happen to pore distribution and pore growth under such a condition - an aspect not yet reported in the literature. Addressing this serves as the second motivation for this work.

In the present study, we revisit the electroporation of a spherical GUV in lieu of a biological cell numerically, in an attempt to understand the response of biological cells to external electric fields, essentially due to the simplicity and tenability of the bilayer membrane constituting the GUV as well as its internal and external environment. In this context, GUVs subjected to pulsed DC fields akin to those used in cell electroporation has attracted significant attention. Strikingly, unusually large prolate, oblate and cylindrical deformations have been observed in spherical GUVs subjected to strong pulsed unipolar DC fields [28] and more recently square and sinusoidal bipolar electric pulses [19]. It is conjectured that the extraordinarily large deformation seen in GUVs is on account of electroporation, including formation of giant pores. It is suggested that the generated pore area manifests as deformation of the spherical GUV. However, no suitable model in GUVs has been suggested to this effect, although few simulations studies have been conducted on simultaneous electroporation-electrodeformation, modelled by the constitutive equations for membrane mechanics. To this end, we validate the recent experimental data of Maoyafikuddin and Thaokar [19]. We primarily emphasize developing an improved electroporation framework that can be able to bridge the research gap by providing mechanistic insight into the formation of giant pores under experimental conditions. For this purpose, we consider electric pulses of unipolar and bipolar types with pulse duration (*T*_*p*_) of 1 ms. Similarly, two charging conditions are studied: (a)*τ*_*c*_ *≪ T*_*p*_ and (b) (a)*τ*_*c*_ *∼ T*_*p*_, to reveal the significance of *τ/T*_*p*_ on the pore arrangement and their evolution. Including the effect of Maxwell’s stress, we observe the pore can grow up to size as large as *∼* 1*µ*m. The present model agrees well with the qualitative trends in temporal evolution of deformation in a porated-vesicle obtained by Maoyafikuddin and Thaokar, thereby serving as a precursor model for predicting porated-vesicle dynamics.

## Electroporation model

### Governing equations

Consider a spherical vesicle of radius *a* and membrane thickness *h*, surrounded by an aqueous medium. An electric pulse of strength *E*_0_, which is spatially uniform, is applied along the axis of symmetry i.e. the z-axis. For simplicity, only the electroporation of the vesicle under the field is considered the associated deformation and flow-field are disregarded in the analysis. The electrostatics of the problem is governed by the electrical conductivities (*s*_*i*_, *s*_*e*_) and permittivities (*ϵ*_*i*_, *ϵ*_*e*_), where the subscript ‘*i* ‘ and ‘*e*’, used for intravesicular and extravesicular fluids, respectively.

The electric potentials associated with each fluid phase obey the Laplace equation, [31]

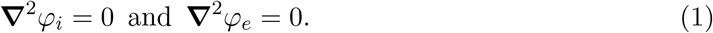

The electric potentials are related to electric field ***E*** as ***E*** = *−****∇****φ*. The presence of the vesicle perturbs the electric potential around it, and the far-field potential converges to the applied potential, i.e., *φ*_*e*_ = *−E*_0_*z*. The applied electric field triggers the accumulation of charges at each side of the membrane over the time scale of,

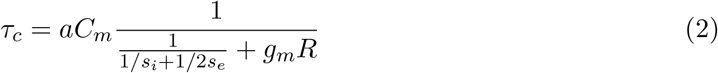

which in the absence of membrane conductance, *g*_*m*_ = 0 is *τ*_*c*_ = *aC*_*m*_ (1*/s*_*i*_ + 1*/*2*s*_*e*_) [12, 31], also known as charging time. Here *C*_*m*_ is the membrane capacitance. In this work we consider *τ*_*c*_ to be the charging time of an unporated vesicle. The charging time *τ*_*c*_ plays a key role in the present investigation. The peripheral charge arrangement induces potential difference at the membrane (*V*_*m*_), also called as TMP, which is written as *V*_*m*_ = *φ*_*i*_ *− φ*_*e*_ at *x ∈ S*, where *S* describes the membrane surface. The current density is continuous across the membrane, which can be expressed as [8, 32, 33]:

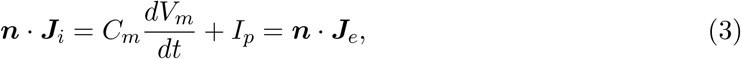

where ***J*** (= *s****E***) is the current density. The first and second terms in the membrane current (Eq. (3)) represent the membrane capacitive current and current density associated with large aqueous pores. The current passing through a single large aqueous pore of radius *r*_*p*_ can be calculated as *i*_*p*,*m*_ = *V*_*m*_*/* (*R*_*p*_ + *R*_*i*_) [36, 17]. Here 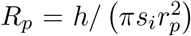 is the pore resistance and 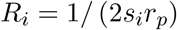 is the input resistance. Accordingly, the net current density due to large pores, as appears in (3) can be calculated as 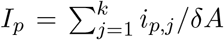, wherein *k* is the total number of pores located in an elementary area *δA*.

Since *V*_*m*_ is established over *τ*_*c*_, the transient electric problem can be explicitly solved using the initial condition, i.e., *V*_*m*_ = *V*_rest_ and *φ*_*i*_ = *φ*_*e*_ = 0 at time *t* = 0. Here, *V*_rest_ is the voltage difference at the membrane in the absence of external electric field, also known as resting potential. The solution to the TMP in the pre-poration period with *I*_*p*_ = 0 can be explicitly derived as

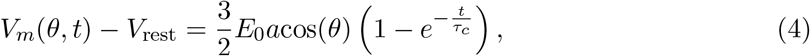

showing a monotonic increase in *V*_*m*_ with time over timescale *τ*_*c*_ for unipolar pulse. Assuming the case of artificially engineered GUVs, we do not consider the contribution of ion-channels, thus, *V*_rest_ = 0 is taken. Similarly, for the bipolar pulse case, the solution for *V*_*m*_ can be obtained as [19]

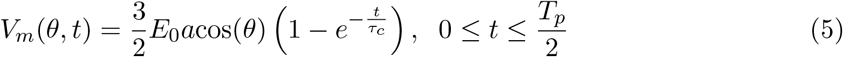

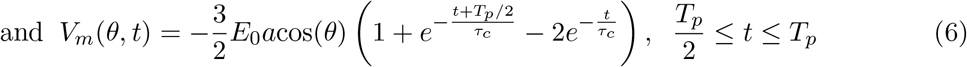

The pore density *N*, which primarily depends on *V*_*m*_ is governed by [23]

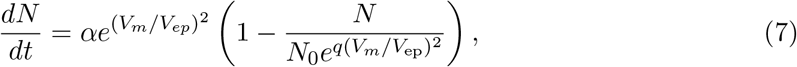

where *N*_0_ is the equilibrium pore density at *V*_*m*_ = 0. In Eq. (6), *α, q* and *V*_ep_ are constants, defined in Table.1 of the earlier paper of Krassowska and Filev [17], and provided in the supplementary material for the sake of brevity. The pores are assumed to form at minimum energy radius, *r*_*m*_ (= 0.8*nm*) [23], and eventually grow under the influence of membrane tension, pore electric energy, and the electric stretching built up in the membrane due to Maxwell’s or electric stress (*τ*_*E*_). While the effects of the former two factors are previously understood, the effect of electric stretching on pore size is exclusively addressed in this work. The electric field develop at the membrane over Maxwell-Wagner time scale, 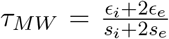, which is much smaller than the membrane charging time, *τ*_*c*_. Thus, the pores must experience the effect of electric stress. Note that irrespective of the type of reported vesicle deformations (*e*.*g*. prolate or oblate), the electric stress leads to membrane stretching thereby promoting pore expansion. The dimensionless electric stress 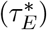 under external uniform electric field, as computed earlier by Vlahvoska *et al*. [39], scales as

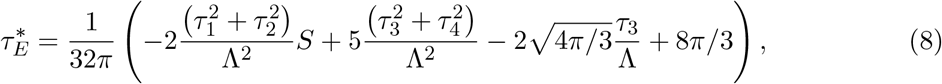

where *τ*_1_ = ((*V*_*m*_ *−* 1)*R* + 1)(*R* + 2) + ((*V*_*m*_ *−* 1)*S* + 1)(*S* + 2)*ω*^2^, *τ*_2_ = *τ*_4_ = *ω*(3 *−* 2*V*_*m*_)(*R − S*), *τ*_3_ = (3 *−* 2*V*_*m*_)((*R* + 2) + (*S* + 2)*ω*^2^) and Λ = (*R* + 2)^2^ + *ω*^2^(*S* + 2)^2^. Here, *R* = *s*_*i*_*/s*_*e*_ and *S* = *ϵ*_*i*_*/ϵ*_*e*_ are conductivity and permittivity ratio, respectively and *ω* represents the frequency of the applied AC electric field. In this work, considering unipolar and bipolar pulsed DC fields, we set *ω* to zero. The corresponding electric tension in the membrane, in its dimensional form, can be approximated as 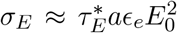. Accordingly, we update the previously formulated pore energy *U* as

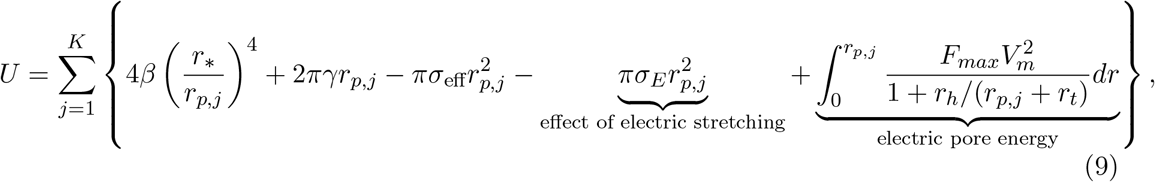

where *r*_*p*,*j*_ is the radius of the *j*^*th*^ pore and *K* is the total number of pores over the vesicle. The pores, which are created at a minimum energy radius *r*_*m*_, evolve in order to minimize the total membrane energy, as

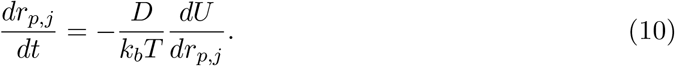

Again, for the various parameters that appeared in Eqs. (8) and (9), please refer to Table 1 provided in the supplementary material. The first, second and third terms on the right-hand side of Eq.(8) are well known, and represent steric repulsion of lipid heads, pore edge energy and membrane tension-effects due to poration, respectively. The fourth term describes the pore energy due to the electrical tension built in the membrane, while the fifth term is the electric force generated at the pore by the transmembrane potential.

### Numerical implementation

The assumption of a spherical vesicle allows us to discretize the computational domain in spherical coordinates (*r, θ*, Φ). Adhering to the assumption of symmetry about the vertical axis, we discretize Eq.1 using the finite difference method. A semicircular computational domain of radius 3*a* is considered, imposing the necessary boundary conditions at the edge of the domain and at the axis of symmetry. A uniform polar grid structure is used to discretize the domain with 100 grids in the angular and 60 grids in radial directions. The obtained grid size (Δ*θ*, Δ*r*) = (*π/*100, *a/*20) is fine enough for the precise estimation of the spatial derivatives near the surface and is ascertained by grid-independence studies.

The initial value problem is solved by initializing the values of the variables *ϕ*_*i*_, *ϕ*_*e*_, *V*_*m*_, *N, I*_*p*_ to zero. At *n* + 1^*th*^ time step, 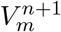 is calculated by applying Euler’s forward method to Eq.3, providing the known value of *I*_*p*_ from the previous time step. The linear set of equations resulting from the spatial discretization of Eq.1 along with the temporal discretization of Eq.3 is solved using iterative methods. Next, considering the stiffness of Eq.5, the midpoint implicit method is utilized to calculate *N*^*n*+1^, using the calculated value of 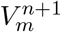. An adaptive time-stepping method is used, which varies the time step varying between 10^*−*9^-10^*−*7^*s* to minimize the computational cost. The time adaptation is incorporated in the following manner. To capture the quick TMP development precisely at the beginning, time step of 10^*−*8^*s* is considered. Eventually, when the poration starts, it leads to sharp increase and decrease in pore density and TMP, respectively; therefore a smaller time-step of 10^*−*9^*s* is taken. In the following stage, i.e., in pore growth, time step of 10^*−*7^*s* is considered. The total number of pores and number of newly launched pores at any angular location *θ* at the *n* + 1*th* time is calculated as 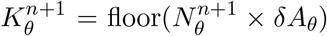 and 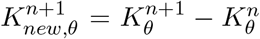, where *δA*_*θ*_ = 2*πa*^2^*sin*(*θ*)Δ*θ* for all *θ* except for *θ* = 0 and *π* where is equal to 2*πa*^2^(1 *− cos*(Δ*θ*)). The total number of pores is symbolized as *K* = (*K*_*θ*_). Note that *δA*_*θ*_ is nonuniform function of *θ*, i.e., it is largest at the equator and smallest at the poles. For such a setup, the model may produce a peculiar error for parametric spaces that results in small pore density all over the membrane. Despite having the highest pore density at the poles, the model may not predict pores at the poles, whereas the nearby locations may contain some pores. This particular error can be eliminated by considering a slightly coarser grid. For this, we consider Δ*θ* = *π/*60 after examining that it negligibly affects the total pore number and TMP.

The 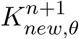 number of new pores are launched at a radius of *r*_*m*_ = 0.8*nm*. To fix their identity, they are assigned with different radii using normal distribution about *r*_*m*_ with a standard deviation of 10^*−*12^*m*. Both the newly launched and existing pores grow following Eq.9, which is solved using the Runge-Kutta method. It is mention-worthy that in Eq.8 the effective membrane tension 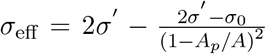 [24] decreases with increase in total pore area 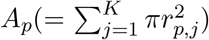, and beyond a certain limit it may attain a negative value, which can not be realized in practice. Thus, the minimum value of *σ*_eff_ is assigned as zero. We realize that the calculation of the force on the pore using the expression 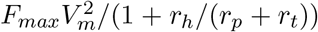 [21] (refer to Eq.8) holds good as long as the pore strictly resides to a single discrete annular region i.e. 2*r*_*p*_ *< a*Δ*θ*. However, for much larger pores, the same can not be used as there can be considerable variation in TMP along its length. As no previous theoretical and numerical understanding has been developed on this subject, the most suitable way of correcting the integrand in the last term of 8 is using area averaging method, written as follows:

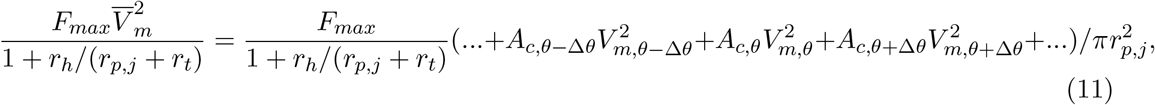

where *A*_*c*,*θ*_ is the projected area of the pore shared by the annulus located at *θ*. Similarly, the current through the pores is modified as

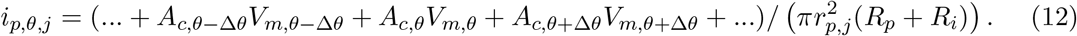

While, ideally, Eq.(9) should be solved for each pore, implementing the same is quite expensive when the total number of pores *K ∼ O*(10^5^), which usually occur for very high *E* and *τ*_*c*_ *<< T*_*p*_. Only for such cases, we adopt an alternative approach that does not sacrifice the accuracy of the solution. In this approach, the set of pores located at a particular *θ* are assumed to have same radius (can also considered as their mean radius), which evolves with time. Accordingly, current density can be calculated as *I*_*p*,*θ*_ = *K*_*θ*_*i*_*p*,*θ*_*/δA*_*θ*_. However, in case of fewer pores, all equations are solved individually for each pores.

The above-described numerical implementations are rigorously validated by comparing our results with the earlier numerical results obtained for the cases of with and without poration (refer to supplementary information). All the simulations are performed using the computing platform MATLAB [20].

## Results and Discussion

### Mechanism of electroporation for different *τ*_*c*_ : unipolar pulse case

The charging time scale *τ*_*c*_ describes the time required for the free charges to accumulate at the membrane surface and depend upon the membrane and fluid electrical properties 2. The value of conductivity ratio (*R* = *s*_*i*_*/s*_*e*_) and radius of the vesicle (*a*) are taken as 2 and 15*µ*m in this work unless explicitly mentioned otherwise. The outer and inner fluid permittivities are the same as that of water. The choice of electric properties is adopted from the recent experimental study of Maoyafikuddin and Thaokar [19].

The effect of *τ*_*c*_ is explained in this section by comparing the cases of *τ*_*c*_ *<< T*_*p*_ and *τ*_*c*_ *∼ T*_*p*_. For *τ*_*c*_ *<< T*_*p*_, TMP develops almost instantaneously, in accordance with Eq. 5. This leads to the immediate pore formation and consequently a sharp increase in pore area on applying electric field, as observed in Fig. 2a. On the contrary, for high *τ*_*c*_ *∼ T*_*p*_(*∼* 1*ms*), pore formation does not occur till time *≈* 0.4 ms as shown in Fig. 2b. However, on the commencement of pore formation, the pore area sharply rises and eventually attains a much higher value than for *τ*_*c*_ *<< T*_*p*_. To analyze this intricacy, the pore density and pore area variations along the polar angle is displayed in the insets of Fig. 2. It is observed that in both cases, the pore density decreases from its maxima at the poles (i.e. *θ* = 0 and *π*) to the equator (*θ* = 0), showing almost a cos(2*θ*) like variation. However, the pore density in the polar regions for *τ*_*c*_ = 0.6*T*_*p*_ is almost an order lower than that of *τ*_*c*_ = 0.01*T*_*p*_. A significant difference is also found in pore area (*A*_*p*_) variation, as represented by contour plots in the insets of Figs. 2a and 2b. An anti-symmetric distribution in TMP and symmetric distribution pore density is observed about the equator. For *τ*_*c*_ = 0.01*T*_*p*_, *A*_*p*_ is minimum at poles and increases monotonically until critical angles are reached. Here, the critical angle (*θ*_*c*_) refers to the angle beyond which a pore does not form. For the parameters considered, the critical angles are *≈* 51^*°*^ and 129^*°*^. For *τ*_*c*_ = 0.6*T*_*p*_, the pore area distribution is almost uniform throughout the poration zone, except in the polar regions, where it is minimal.

**Figure 1:**
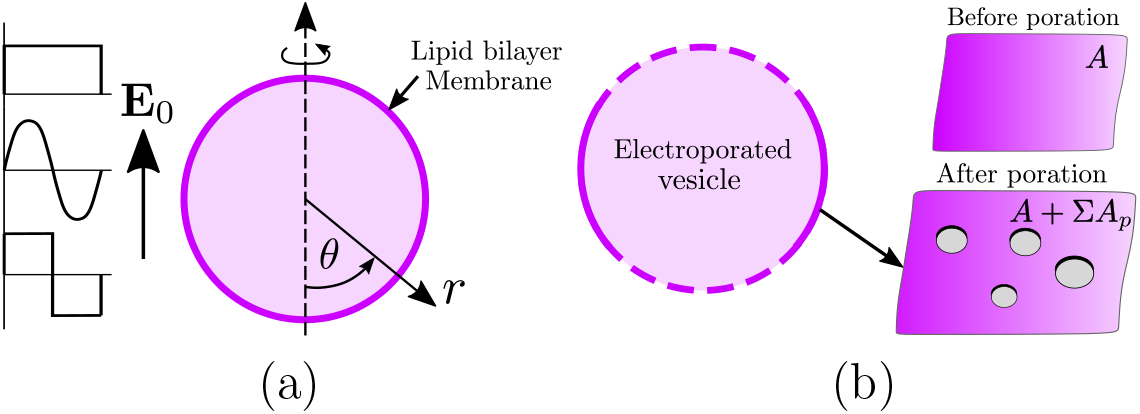
Schematic representing (a) a spherical vesicle under electric pulses before poration and (b) after poration.

**Figure 2:**
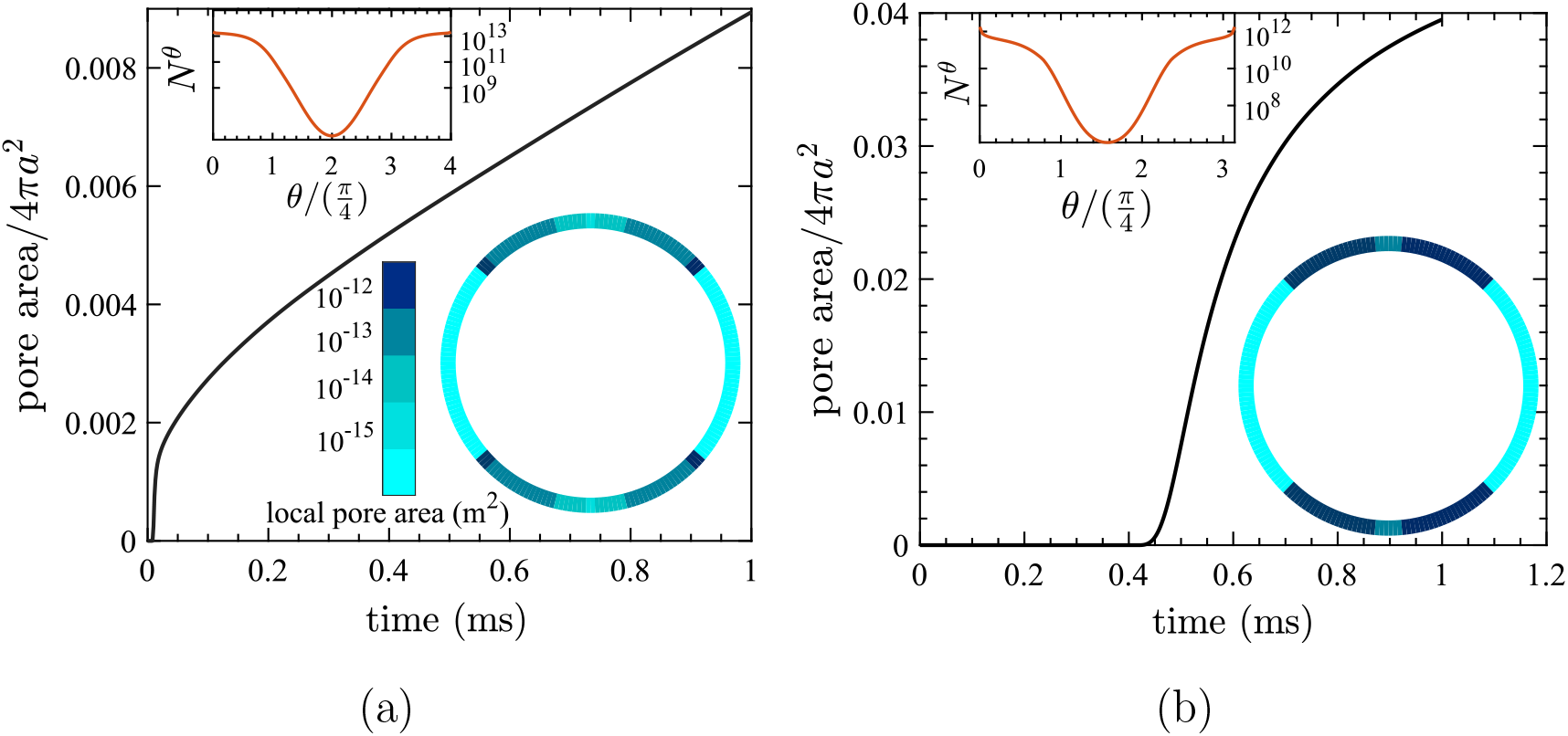
(a) and (b) present the time variation in total pore area for *τ*_*c*_*/T*_*p*_ =0.01 and 0.6, respectively. The pore area distributions along the surface are represented by colour contours. The pore density *N*^*θ*^ variation along the surface are provided as insets. The considered electric field is a unipolar pulse of strength 100 kV/m

The variations in pore density and pore area, as illustrated in Fig. 2, project two surprising features. Firstly, although the pore density at the poles is significantly high, the local pore area is quite small. Secondly, even though the increase in *τ*_*c*_ results in fewer pores, the total pore area is larger. This can be understood from the transient variations in maximum and mean pore radius i.e. *r*_*p*,*max*_(*t*) and *r*_*p*,*m*_(*t*), for *τ*_*c*_*/T*_*p*_=0.01 shown in Fig 3a and 3b, respectively. It is apparent that the radii of the pores near *θ* = 0 remain in the *O*(10)*nm*. Variation in pore radii with *θ* is small up to *θ ≈* 26^*°*^. The alterations in the pore radii are large around *θ* = 45^*°*^, 135^*°*^, albeit in a small range of *θ*, with pores of *O*(10^2^)*nm*. In fact, in the vicinity of the critical angles, *≈* 51^*°*^ and 129^*°*^, pores can even grow up to a few *µm* in size. Importantly, the growth of these set of pores around the critical angles, contributes to the continuous variation in the pore area, since the pores at other locations on the surface, quickly reach their steady state. Fig. 3c compares the influence of pore electric energy and membrane electric tension on the largest pore. It is clear that the contributions of the latter can be as significant as the former towards the later stage when the pores have grown big. This accounts for the fact that the energy due to membrane electric tension depends on the size of the pore, such that its influence on the growth of tiny pores is negligible. This indicates that the initiation of the large pores is predominantly caused by the pore electric energy, while once the pores have grown in size, both the electric energy and the electric tension lead to further pore growth.

**Figure 3:**
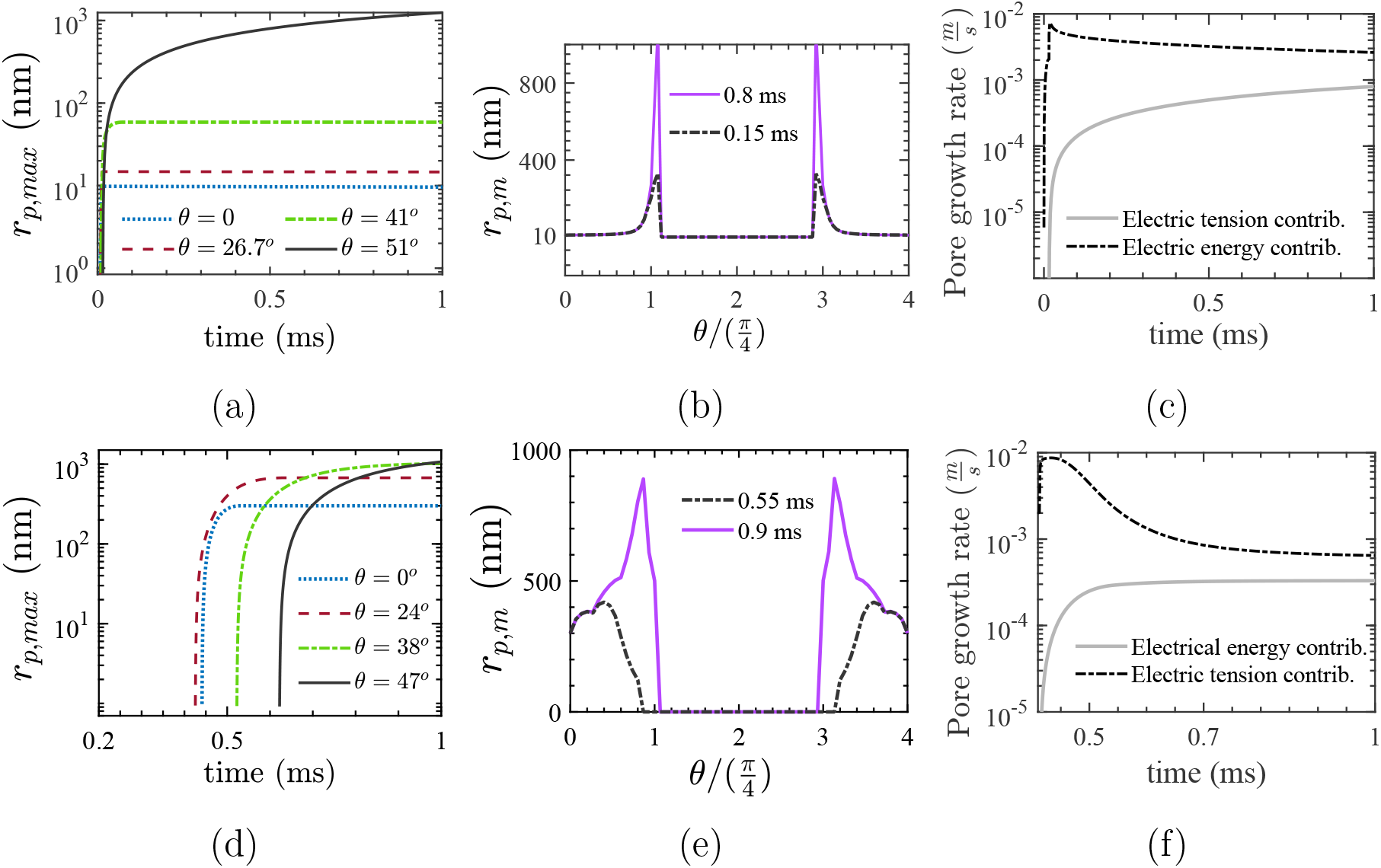
(a and b) shows time variations of maximum pore radii at different angles for *τ*_*c*_ = 0.01*T*_*p*_ and 0.6*T*_*p*_, respectively. (b and e) shows the distribution of mean pore radii along the surface, (c and f) shows the transient evolution of different pore-driving energies. The considered electric field is a unipolar pulse of strength 100 kV/m.

Fig. 3d and 3e depict the transient variations in maximum and mean pore radius i.e. *r*_*p*,*max*_(*t*) and *r*_*p*,*m*_(*t*), for *τ*_*c*_ = 0.6*T*_*p*_, respectively. In this case, the pores near the poles are found to be an order larger than those found for *τ*_*c*_ = 0.01*T*_*p*_. On the remaining part of the membrane, excluding the no-pore formation zone, pores are found to be of *O*(1)*µm*, leading to significantly more pore area, albeit an order less in number than the *τ*_*c*_ = 0.01*T*_*p*_ case (Inset Figure **??**3a,b). Furthermore, as opposed to the previous case, here, over a majority portion of the membrane, pores exhibit continuous growth, increasing the rate of pore area variation. Fig 3f shows the influence of pore electric energy and membrane electric tension on the pore located at the critical angle (47^*°*^ in this case), describing their similar roles on pore growth as shown in Fig. 3c. Together Figs. 3c and 3f show that irrespective of *τ*_*c*_ the pore electric energy reduces with the increase in pore size.

To understand the qualitatively different variations in pore distribution and pore radii shown in Figs. 2 and 3, the TMP variations with time are provided in Figs. 4a and 4b at the pole and critical angles for *τ*_*c*_*/T*_*P*_ = 0.01 and 0.6, respectively. The dashed lines represent the temporal variations in TMP and act as reference. On applying the electric field, the membrane charging gradually takes place, developing TMP. When the poration is ignored, the TMP monotonically increases and attains a steady state as described by Eq. 5. In addition, the TMP gradually decreases from its maxima at the poles to its minima (*V*_*m*_ = 0) at the equator, exhibiting a cos *θ* variation, as suggested by the Schwan equation. However, onset and progress of poration, significantly alter these trends. The TMP follows Eq. 5 as long as it is below a threshold limit. At a given angular location, if *V*_*m*_(*θ*) crosses a certain threshold, the pores start to form. In our study, the threshold limits are found to typically fall between 0.8-0.9 V. Consideration of uniform angular discretization in our model generates nonuniform discrete surface areas. Therefore, for a given *N* (*V*_*m*_), slight variations in pore numbers (= *floor*(*NA*_*θ*_)) can be obtained along the membrane (inset of Fig. 2). This makes it challenging to ascertain the exact value of threshold TMP.

**Figure 4:**
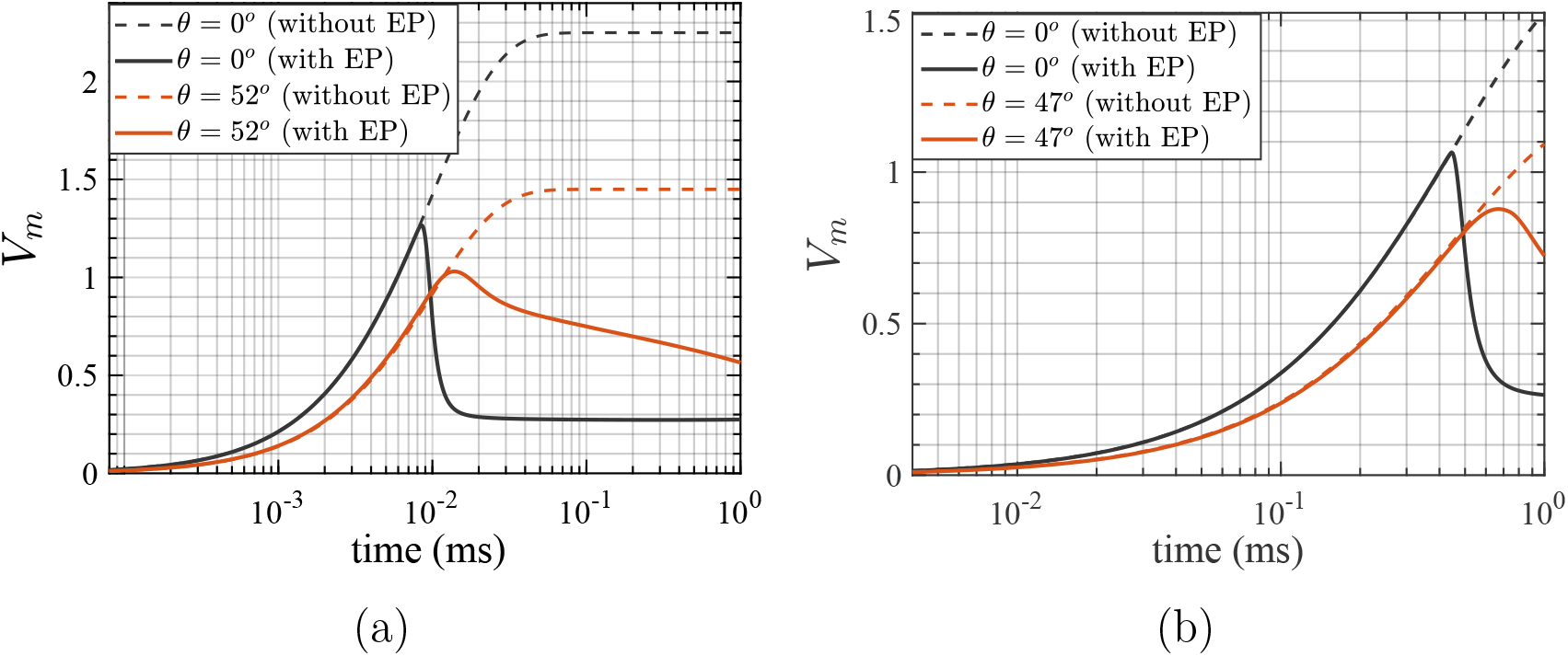
(a and b) show the transient transmembrane potential variations at poles and at critical angles for *τ*_*c*_ = 0.01*T*_*p*_ and 0.6*T*_*p*_, respectively. The critical angles for *τ*_*c*_ = 0.01*T*_*p*_ and 0.6*T*_*p*_ are 51^*°*^ and 47^*°*^, respectively. The considered electric field is a unipolar pulse of strength 100 kV/m

Fig. 4a indicates that the poration initiates first at the poles and then gradually advances towards the equatorial region (See *V*_*m*_(*t*) at *θ* = 0, *θ*_*c*_). The membrane, which behaves as a perfect capacitor before poration, exhibits a precipitous drop in TMP with time at *θ* = 0 shown in Fig. 4a due to formation of a large number of very small pores. The drop in TMP is essentially due to transformation of the perfect capacitor-like membrane to a leaky capacitor, on account of generation of membrane conductance due to poration and admittance of a leakage current. This is also corroborated by the small pore size (Fig. 3b) and corresponding smaller pore area despite a large number of pores at the poles (see inset of Fig. 2a). The high conductance at poles leads to temporal plateauing of TMP (displacement current of the membrane is vanishingly small) due to local shorting of the membrane, which results in the current due to membrane conductance, balancing the ohmic current brought into the membrane by the outer and inner fluids, thereby correspondingly leading to temporal saturation of the number of pores and pore growth (Also see Figure 3a), aided by the very low value of TMP. At *θ* = 52^*°*^, *V*_*m*_(*θ*) attains the threshold TMP limit at longer times, thereby delaying poration (15 *µs* for *θ* = 52^*°*^ vs 9*µs* for poles). The gradual increase in TMP at *V*_*m*_(*θ* = 52^*°*^), essentially due to the cos *θ* factor in the Schwan equation, leads to formation of very small number of pores of small radius, and hence much smaller membrane conductance as compared to that at the poles. Thus, after poration, when the potential drops at *V*_*m*_(*θ* = 52^*°*^), the drop in potential is much gradual due to the low membrane conductance, and the membrane discharge happens very slowly, almost over 100*µs*, aiding pore growth due to a still significant value of *TMP*. In this process, the TMP decreases essentially due to a gradual increase in membrane conductance such that the membrane ohmic current balances the ohmic current due to the intra and extracellular fluids. The slow change in TMP leads to a large growth in pore size and thereby pore area (Figures Inset 2a,3a,b).

The physical quantities *V*_*m*_(*θ*) and *r*_*p*_(*θ*) are coupled in two ways. The pore growth phase, which is preceded by pore formation, is governed by electric energy and membrane electric tension, as mentioned earlier. It can be examined that the membrane electric tension is a weak function of *V*_*m*_, depending on the applied field, whereas pore electric energy scales as 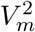. Accordingly, small *V*_*m*_(*θ* = 0) post poration leads to insignificant pore growth, which is eventually arrested by the edge tension. As a result, the pores at *θ* = 0 stop growing resulting in the plateauing of current density *I*_*p*_(*θ* = 0) and *V*_*m*_(*θ* = 0). Conversely, at *θ* = 52^*°*^, the continuous pore growth and concomitant increase in current reduces the TMP. This explains the temporal weakening of the pore electric energies shown in Figs. 3c and 3f.

Figure 4b presents the TMP variation for *τ*_*c*_ = 0.6*T*_*p*_, where transients are quite slow both during the charging and the poration phase. A closer look into Figs. 4b and 4a reveals that the peak value of *V*_*m*_ at a *θ* = 0 in the case of *τ*_*c*_ = 0.6*T*_*p*_ is relatively lower than that of *τ*_*c*_ = 0.01*T*_*p*_, on account of gradual development of membrane conductance owing to higher *τ*_*c*_, thereby explaining the lower pore density as shown in Fig. 2b as compared to *τ*_*c*_ = 0.01*T*_*p*_. Surprisingly, though both for *τ*_*c*_ = 0.01*T*_*p*_ and *τ*_*c*_ = 0.6*T*_*p*_, the TMP at *θ* = 0 comes down to *≈* 0.3V and the pores observed in the latter case are much bigger at *θ* = 0 due to slower development of TMP. This also results in a higher mean pore radius across a majority of the angular locations in the latter case despite the delay in pore formation.

### Effect of bipolar pulse

Based on the understanding developed in the preceeding section, we now investigate the effect of bipolar pulse which is also widely employed in several medical procedures, on GUV electroporation. The mathematical expression for the applied bipolar pulse is as follows: *E* = *E*_0_ for 0 *≤ t ≤ T*_*p*_*/*2, *E* = *−E*_0_ for *T*_*p*_*/*2 *≤ t ≤ T*_*p*_. Figure 5a shows the transient pore area variation for different charging times. It is observed that for *τ*_*c*_ = 0.01*T*_*p*_, the pore area variation for bipolar and unipolar pulse are almost identical. Increasing *τ*_*c*_ delays the pore formation as well as results in lesser pore density (shown in the inset), as discussed before. The number of pores obtained for *τ*_*c*_*/T*_*p*_ =0.01, 0.1, and 0.6 are *≈* 5100, 650, and 100, respectively. Despite lesser pores, for both, *τ*_*c*_ = 0.1*T*_*p*_ and 0.6*T*_*p*_, the pore area remarkably rises rapidly, followed by a more modest increase. For high *τ*_*c*_, the pore area for bipolar pulse drastically differs from the same found for unipolar pulse. It is mention-worthy that the pore area for *τ*_*c*_ = 0.6*T*_*p*_ is shown to eventually attain a steady value. However, in reality, our model initially predicts decrease in pore area after reaching a certain value owing to pore closure. This decrease in pore area, a manifestation of high *τ*_*c*_, is artificial ignored in our numerical code to support the fact that the vesicle relaxations usually occur over a time scale of *O*(10) *−O*(100)*ms*, as experimentally demonstrated by Yu *et al*. [40]. Under the considered pulse duration i.e. 1*ms*, therefore, the vesicles can be assumed as quasi-stable, potentially resisting any surface area modulations induced by large-pore closure. In fact, a number of studies can be found in the literature suggesting that the pore closure time can typically vary between *O*(1) *− O*(100)*ms* [15, 9, 30], though the exact reason still remains as an open question for the research community. Along with these reasons one may need to consider the electric tension effect, which counters the pore closure to some extent. Undoubtedly, the pore closure can not be realized in the considered pulse duration and as this issue can not be resolved using the existing electroporation models. Therefore, we artificially implant pore closure termination in our simulations, upon which, our model effectively c orroborates w ith the previous e xperimental fi ndings [1 9]. (refer to the last section for detailed discussion).

**Figure 5:**
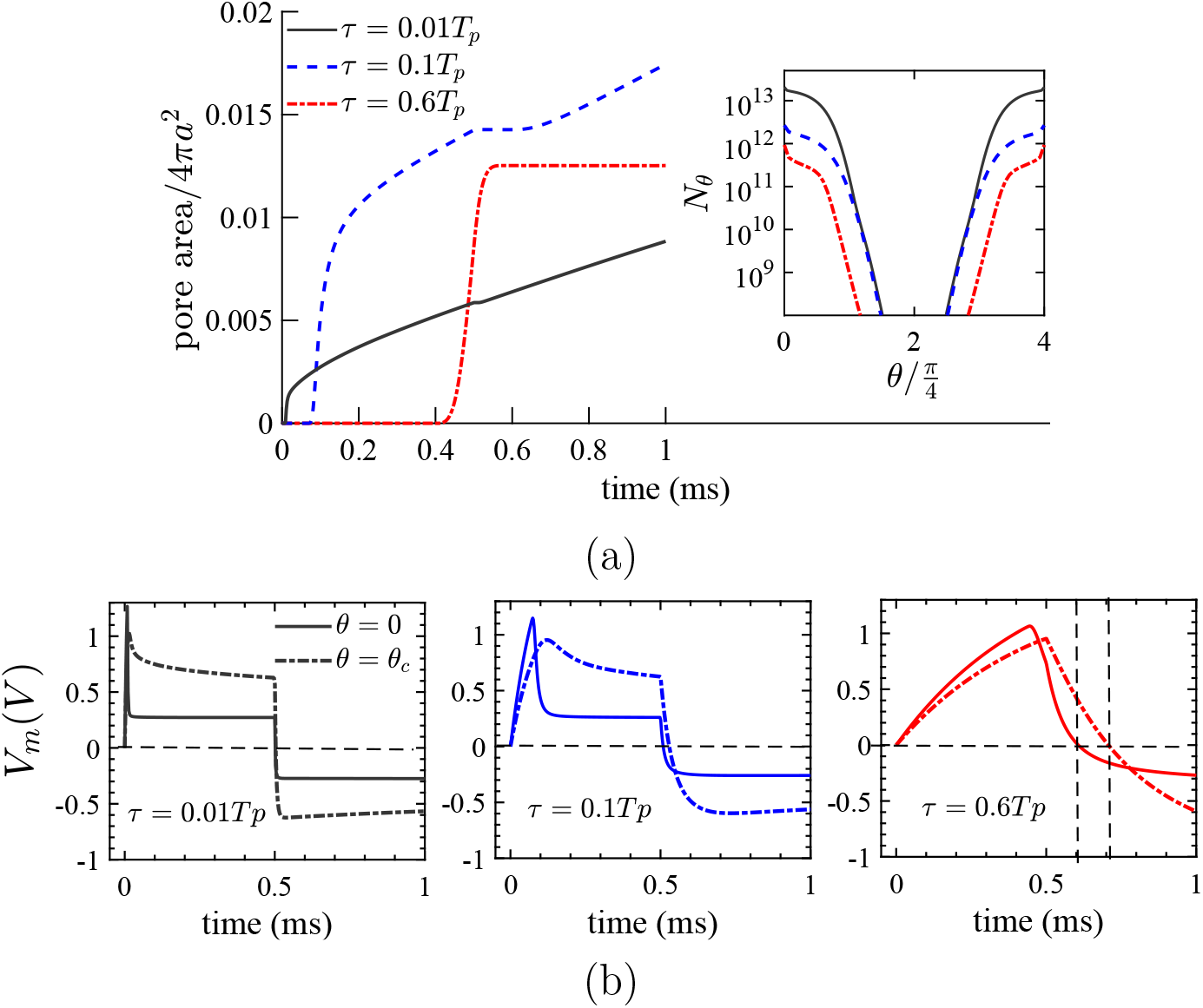
(a) Transient surface area variation under bipolar pulse for different charging time. The inset of (a) shows the respective pore density as a function of polar angle. (b) Transmembrane potential time-variations at poles and polar angles for different charging time. The strength of considered bipolar pulse is 100 kV/m

From Fig. 5a suggests that the pore area sensitively depends on the ratio of *τ*_*c*_*/T* appears to admit an optimal ratio for a bipolar pulse, where it is maximum. To gain insight, we analyze the TMP variation along the surface for different *τ* _*c*_ i n F ig. 5 b. For 0 *≤ t ≤ T*_*p*_*/*2, the TMP variations in the pre-poration period is the same as that of a unipolar pulse see Eq. 4 and 5.

The poration for *τ*_*c*_ = 0.01*T*_*p*_ occurs much before the switching of polarity. The short charging time then leads to an instantaneous reversal of TMP throughout the membrane on reversing of the polarity of the bipolar pulse, whereby the magnitude of the TMP is unaltered. Thus, the pores do not sense any significant d iscontinuity (reduction) i n e lectric e nergy a nd h ence the pore growth for bipolar pulse is obtained to be the same as an unipolar pulse of same *T*_*p*_. In the extreme, for a very high charging time *τ*_*c*_ = 0.6*T*_*p*_, due to delayed pore creation, the pores have very little time to grow before the phase reversal begins at *t* = 0.5ms. The reversal of TMP occurs slowly over the long charging time *τ*_*c*_, and the relaxation for an extended period, wherein, the TMP goes from a positive to a negative value, through a state of 0 TMP, results in the TMP all over the membrane remaining consistently low (|*V*_*m*_| *<* 0.25V) as displayed in Fig. 5b. The pores hardly grow in this low-energy state, and in some cases, can even exhibit initiation of pore closure, especially in large pores, influenced by strong edge tension forces. Consequently, as compared to an equivalent unipolar pulse, a much-reduced pore area is observed. However, this obtained pore area is still much higher than that obtained for *τ*_*c*_ = 0.01.*T*_*p*_, since the majority of the pores *τ*_*c*_ = 0.6*T*_*p*_ are quite large, as illustrated before for the unipolar pulse case.

For *τ*_*c*_ = 0.1*T*_*p*_, the pore growth is larger than *τ*_*c*_ = 0.01*T*_*p*_, and faster than *τ*_*c*_ = 0.6*T*_*p*_ leading to larger pore area before polarity reversal. On polarity reversal of the applied pulse, the *V*_*m*_ changes sign over timescales, slower than that for *τ*_*c*_ = 0.01*T*_*p*_, thereby reaching lower TMP than an equivalent unipolar pulse, but faster than *τ*_*c*_ = 0.6*T*_*p*_, consequently spending lesser time around *V*_*m*_ = 0. Thus, the pore area for *t*_*c*_ = 0.1*T*_*p*_, seems to be greater than that for 0.01 and 0.06 *T*_*p*_.

### Effect of electric field strength

The charging time not only regulates the pore number and their size but also regulates the angular extents of the prorated regions, as displayed in Fig. 6. To illuminate the same, we plot *θ*_*c*_ vs. *τ*_*c*_*/T*_*p*_ for bipolar pulses of strength 50, 75, 100, and 125 kV/m. For |*E*| = 125 kV/m, *θ*_*c*_ is almost insensitive to the change in charging time, and it remains constant at *θ*_*c*_=51^*°*^ up to *τ*_*c*_ *≤* 0.6*T*_*p*_ before slightly decreasing when *τ*_*c*_ increases to 0.7*T*_*p*_. For weaker pulses, this decline in *θ*_*c*_ starts at much lower values of *τ*_*c*_. However, such variations are less pronounced for unipolar pulses [17].

**Figure 6:**
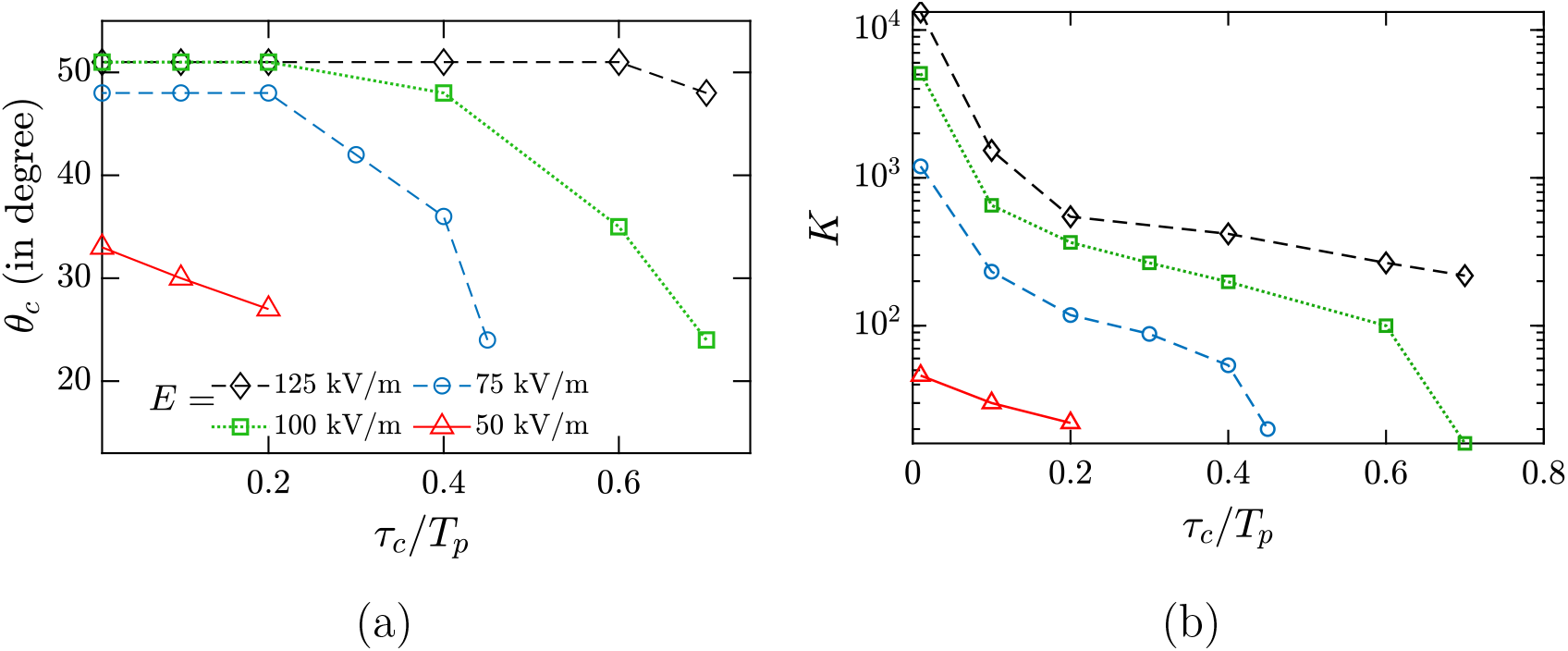
(a) Critical angle and (b) total pore number (*K*) as a function of charging time for bipolar pulses of different strengths.

The decrease in critical angle with electric field is equivalent to the narrowing of the poration zone and affects the total pore number (*K*), as delineated in Fig. 7b. Increasing charging time adversely affects *K*, while an increase in pulse strength enhances *K* by generating a higher TMP in the pre-poration phase. It is apparent that *K* is a strong function of |*E*| as increasing |*E*| from 50 kV/m to 125 kV/m can increase the *K* by almost two orders.

**Figure 7:**
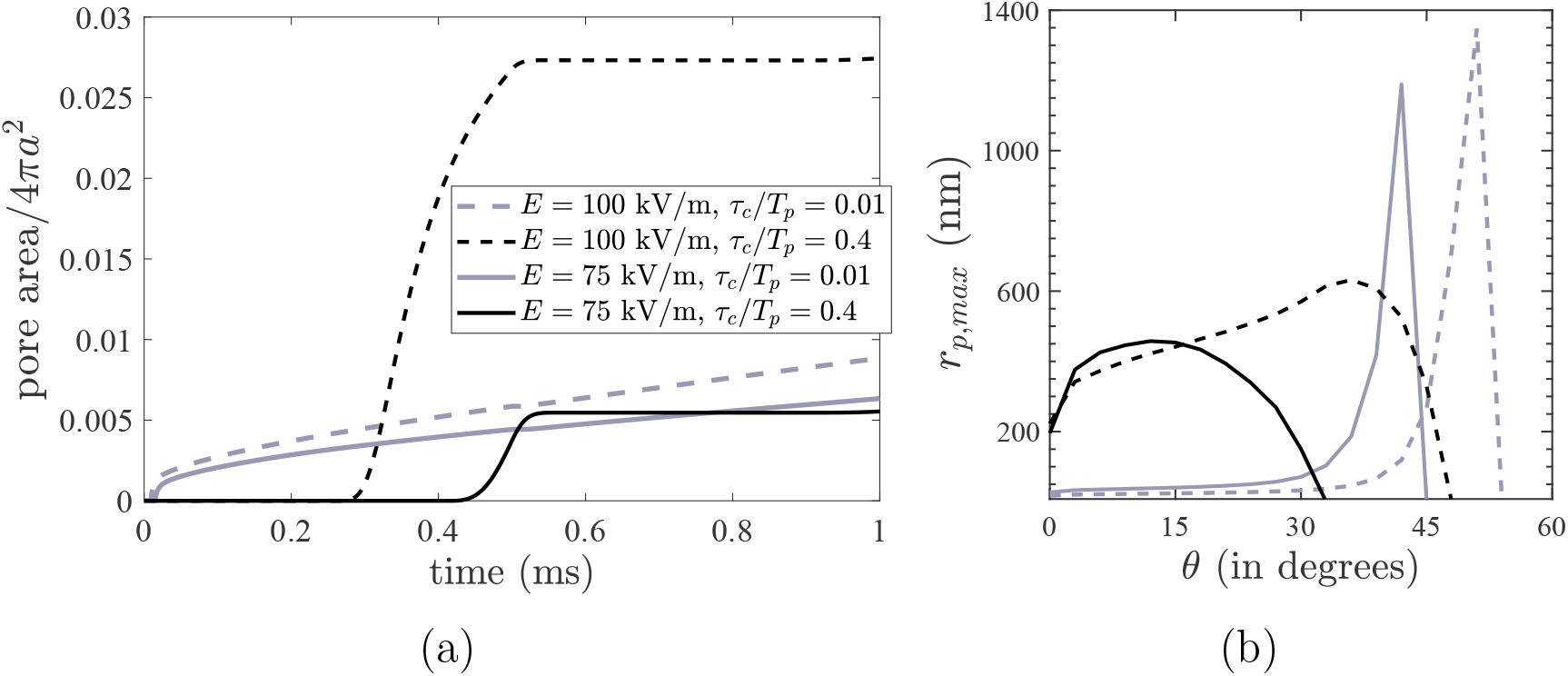
(a) Effect o f bipolar pulse s trength o n total p ore area f or different charging time. (b) Maximum pore radii variation with polar angle for different p ulse s trength a nd charging time.

Further, an increase in the electric field |*E*| affects the pore area as displayed in Fig. 7a and is extremely sensitive to *τ*_*c*_. For *τ*_*c*_ = 0.01*T*_*p*_, a very small increment in pore area is obtained on increasing electric field strength, say from 75 kV/m to 100 kV/m, despite one order of increase in *K* (see Fig. 6b. This is attributed to the fact that at small *τ*_*c*_, an increase in |*E*| leads to more pores but of smaller radius, except around critical angles Fig. 7b. The smaller radius is because of a larger drop in TMP due after operation at greater |*E*| fields, causing lesser pore growth. Thereafter, the increase in pore area is mainly driven by the growth of large pores around critical angles.

Notably, while for *τ*_*c*_ = 0.01*T*_*p*_, the |*E*| has an insignificant effect on the transients of the pore area, it plays a vital role in modulating the transients for systems with high *τ*_*c*_. Referring to Fig. 7a for cases of *τ*_*c*_ = 0.4*T*_*p*_, it can be observed that for |*E*| = 100 kV/m, the initiation of pore formation and growth happens much earlier, reflecting in the early rise in pore area compared to |*E*| = 75 kV/m. However, for both the values of |*E*|, the pore growth ceases at almost the same time, at 0.5ms in Fig. 7a when the polarity of the applied bipolar pulse changes. This means the pore growth spans for a longer period for larger |*E*|. The early initiation of pore growth for larger *E*, thus results in larger pore area. Additionally, contrary to low *τ*_*c*_, in the case of high *τ*_*c*_, at higher |*E*|, larger pores are created on nearly the entire porated region that grow up to *O*(100)*nm*, thereby sensing much stronger electrical stretching. Therefore, although strong electric fields result in lower TMP, this decremental effect is compensated by a greater increment caused by pore stretching at higher fields. Accordingly, our model predicts the formation of larger pores at higher |*E*|, in contradiction to the findings of Krassowska and Filev [17], who did not consider electric stretching in their model.

### Estimation of vesicle surface area using present electroporation model

Lastly, to assess the performance of our model and to establish a relation between the transients in electroporation and electrodeformation, we compare our numerical results with the recent experimental results of Maoyafikuddin and Thaokar [19]. In particular, we compare the transient surface area variation of a vesicle as surface area is the most relevant parameter for this study, which is mutually controlled by electrodeformation and electroporation. Though our model is designed to predict poration in a spherical vesicle, it can still be used for predicting the temporal variation of vesicle deformation. The main assumptions here are that the vesicle deformation does not alter the poration characteristics and the large deformations seen in experiments is predominantly a consequence of excess area created through electroporation and the contribution of the inherent excess area as well as due to the unfurling of wiggles in the membrane is negligible. This is much justified, s pecifically fo r th e ex perimental results compared in Fig. 8, where the charging time for the experimental properties is of *O*(*µs*), which implies that the pore formation initiates while sphericity of vesicle shape remain almost intact. The latter drives the pore growth via electric stretching of the membrane. Thus, the surface enlargement of a porated membrane must be attributed to the combined effects of electric stretching and pore growth. The enlargement of the membrane area due to poration translates into extra deformation or higher shape aspect ratio, as observed by Maoyafikuddin and Thaokar [19]. It can be inferred that the additional deformation caused by poration is likely to follow the same transient path as that induced by electric stretching, which allows us to conduct the present comparative analysis. While the experimental deformation data are provided in terms of shape aspect ratio, we compute the respective values of the surface area assuming a prolate shape for the vesicle. In the absence any established formula, the surface area of a porated-only vesicle is determined by the simple summation of the total pore area and initial surface area.

**Figure 8:**
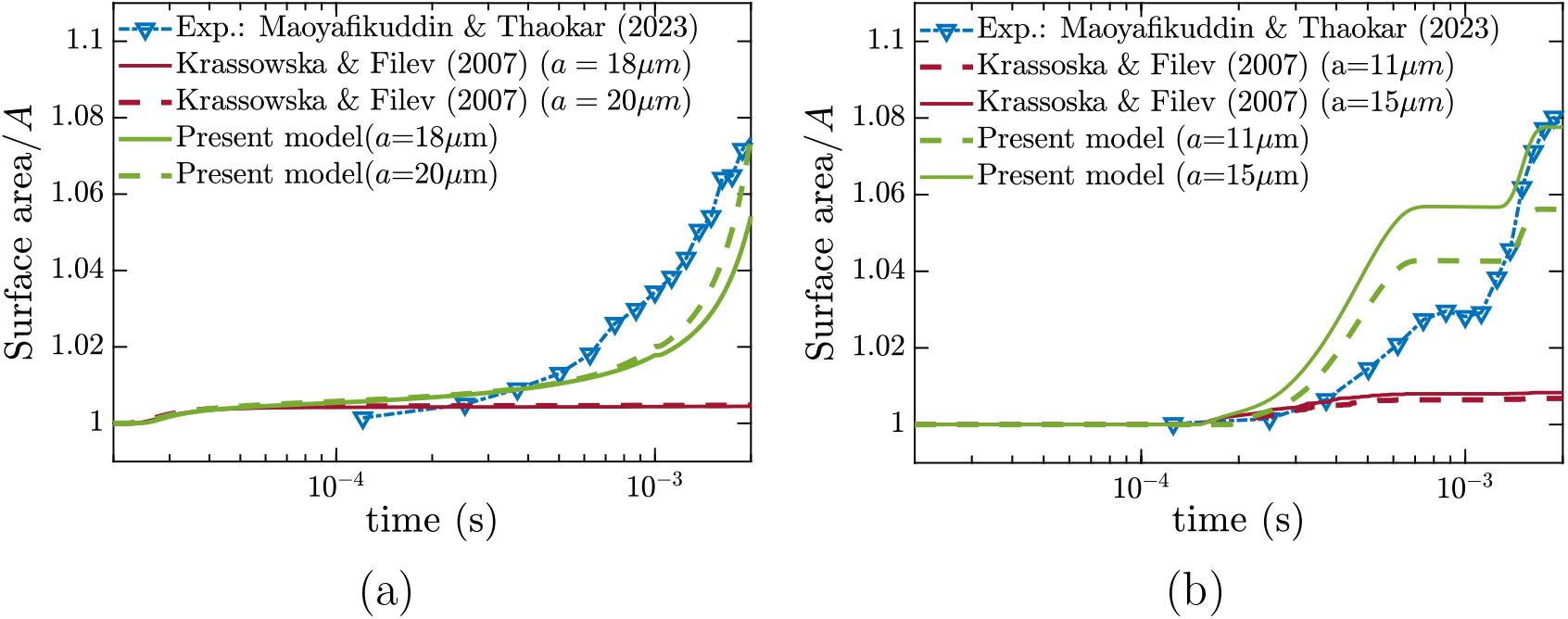
Evolution in surface area of vesicles under (a) bipolar pulse and (b) sinusoidal pulse of strength 100 kV/m.

Consider a vesicle subjected to a square wave bipolar pulse as in the experiments of Maoyafikuddin and Thaokar [19], with experimental conditions as follows: *s*_*e*_ = 18 *×* 10^*−*4^*S/m, s*_*i*_ = 2*s*_*e*_ and *ϵ*_*e*_ = *ϵ*_*i*_ = 80*ϵ*_0_. Fig. 8a illustrates the evolution in surface area under bipolar or square wave pulse of time period 2 ms. The analysis using a one fixed value of the radius of a vesicle for establishing the said analysis is rendered difficult, since the radii of the vesicles employed in the experiments of Maoyafikuddin and Thaokar [19] typically varied between 10 *−* 20*µm*. For this reason, we compare our numerical results considering different radii, as shown in Fig. 8 with the experimental results of [19]. It can be observed that both our numerical and the experiments in Maoyafikuddin and Thaokar show negligible variation in vesicle area up to *t ≈*0.2 ms. At *t ≈*0.5 ms, the surface area starts to rise gradually, and can be attributed to electrodeformation-poration. Although considerably simpler, the model captures the qualitative features very well due to the inclusion of electric stretching effect. On the other hand, the existing model of Krassowska and Filev [17] is unable to predict the data even qualitatively. Thus the stretching term seems to be critical to the observation.

The comparative study is further extended to sinusoidal pulse case, illustrated in Fig. 8b, for all other electrical parameters remaining the same. Although qualitatively in agreement with the experimental results like the previous case, here, the numerical code considerably overpredicts the surface area observed in experiments. However, with the decrease in vesicle radius, the discrepancies are significantly minimized. One of the reasons behind the discrepancy is electrodeformation and electroporation are coupled phenomena as each of these phenomena can individually modulate the transmembrane potential and, thereby, can influence the other. As in the experiments, the vesicles undergo cylindrical deformations; only the end region experience normal electric field. As the transmembrane potential is largely governed by the normal electric field (refer to Eq. 3), except in the polar region it becomes zero everywhere. This constrains the pores only to polar regions, reducing the total pore area. Additionally, due to the unavailability of the actual membrane properties, we adopt the data provided by Smith *et al*.[36] in our study (refer to Table 1 of their paper), which might be different, and one of the main reason behind the discrepancies, apart from ignoring effect of shape change of vesicle on the electrostatics. The larger pore area reached in sinusoidal pulse as compared to square wave pulse can be attributed to slow variation of TMP in the sinusoidal case, leading to fewer larger pores. This may lead to intra-vesicular fluid loss and consequent reduction in vesicle deformation, explaining which is beyond the scope of present model. However, the key limitation of our model is that its reliability for cases where charging time is of *O*(*ms*) as in such scenarios pore formations can be seen to occur much after the deformation violating the assumption of initially spherical shape.

## Conclusion

In summary, a novel numerical approach is proposed to examine the electroporative response of a GUV to milli-second long unipolar and bipolar DC pulses. The highlight of the model is the accounting of electrodeformation-induced effect on pore dynamics. The effect of membrane charging time on TMP, pore density and pore growth is studied. It is found that increasing the charging time leads to slows down the temporal variation in TMP, accordingly takes higher time to attain the threshold value required for initiating poration. Besides, it also decreases the pore density. The poration occurs only up to a certain angular location from the poles on the membrane, named as critical angle. For charging time much smaller than 1 ms, near the critical angles, the pore growths are massive, transforming the pores to giant ones. However, for charging time *∼ O*(1)*ms*, almost all the pores except that restricted to the polar region grow into giant pores. Therefore, the latter case produces total pore area even with less number of pores. For weak electric pulses, it is observed that beyond a certain charging time, the poration can be completely avoided as the TMP may not reach the threshold value before the termination of electric field. The present numerical results aligns well with the earlier experimental results on qualitative grounds.

